# CREation of an expanded plant memory gene circuit toolkit

**DOI:** 10.1101/2025.10.28.684748

**Authors:** Patrick Gong, Adil Khan, Florence Ly, Jia Yuan Zhu, Gabrielle Herring, Harish Jadhav, Christian Pflueger, James P B Lloyd, Ryan Lister

## Abstract

Genetic circuits rely on modular, well-characterized genetic parts to achieve predictable cellular behaviour. Despite their widespread adoption, biological parts are complex and when used in a new molecular context can result in unexpected outcomes. Site-specific recombinases have been widely used due to their unique ability to induce precise, irreversible changes to a DNA sequence, making them ideal tools for memory logic operations. However, unexpected context-specific failures of even the most widely used recombinase, Cre, has limited the expansion and complexity of synthetic gene circuits in diverse species. Here, through a systematic analysis of Cre failure in plant gene circuits, we uncovered multiple unexpected post-recombination interactions within the transcriptional unit between the promoter region, recombinase, and their cognate recognition sites. These significantly inhibit transcriptional activity, preventing circuit functionality. Specifically, post-recombination Cre recombinase exhibits an inhibitory property by binding to *lox* sites, with the remaining *lox* site repressing transcription. By thoroughly characterizing these dynamics, we restored Cre functionality and expanded the plant-based recombinase toolkit for Cre-related recombinases, developing combinations of recombinases and cognate sites with different activation levels for constructing logic gates with stable memory functions. We further developed multiple functional split-recombinase systems for simultaneous logic using a range of dimerization domains and constructed complex logic gates, including 3-input and 6-input AND gates. Together, this exploration of component context dependency in genetic circuits, even with commonly used parts like the Cre/*lox* system, has implications extending beyond plant systems, and significantly expands plant memory circuit capabilities.

## Introduction

Synthetic gene circuits constructed from modular genetic components, akin to electronic circuits that perform complex computational operations, have become indispensable to synthetic biology for the precise control of cellular functions. Yet, despite substantial progress in the characterization and development of standardized genetic parts for transcriptional regulation, reliably achieving predictable and robust circuit function remains a significant challenge in biology (1, 2). The inherent complexity of coordinating multiple regulatory factors and elements within a biological system can often result in the emergence of unexpected interactions or context-dependent behaviors when introduced into non-native environments (2–5).

Site-specific DNA recombinases are well-established and characterized genetic tools for recombining specific target sequences within a genome through the removal, insertion, or reversal of intervening DNA sequences (6). These enzymes typically recognize and bind to pairs of short identical target sequences, resulting in double-stranded breaks and subsequent rejoining of DNA to form recombinant products (7). Widely employed for the generation of conditional knockouts (8, 9) and regulating gene expression (10–12), recombinases, such as Cre and Flp, form a versatile toolkit for precise genetic manipulation across diverse organisms (12–14). The distinct ability of these proteins to irreversibly modify DNA enables their potential to perform binary biological operations and the construction of memory-based gene circuits (15, 16). Such recombinase-based circuits have proven popular and effective for programming customized decision-making processes in bacteria (17–19), mammalian cells (15, 20, 21), and plants (16, 21–23). In these studies, the circuits typically use recombinase recognition sites to flank critical sequences in the transcriptional unit, such as promoters, coding sequences, and terminators. The recombinase then catalyzes a recombination event between the two recognition sites, leading to the inversion or excision of the intervening sequence, depending on the orientation of the core recognition sequences, and consequently the transcriptional activation or repression of the target gene. Multiple sets of orthogonal recombinases and their cognate recognition sites could therefore be used to engineer more sophisticated logic operations and enable the construction of increasingly intricate genetic pathways. To date, simple one- and two-input Boolean logic gates as well as a range of more advanced computations, such as a four-input logic operation (19), a three-input-two-output half adder-subtractor (15), and a six-input-one-output Boolean logic look-up table (15) have been constructed using recombinase enzymes.

Despite these achievements, substantial challenges persist with the implementation of recombinase-based circuits. Even well-characterized recombinases can fail to produce the predicted outcome in various contexts. For example, while both Flp and B3 recombinases effectively activated gene expression in a YES gate, B3 exhibited reduced excision efficiency in the initial NOT gate design, resulting in a failed circuit in *Arabidopsis thaliana* (common name: Arabidopsis) cells (16). Similarly, the Cre-mediated excision-based YES gate performed poorly compared to the inversion-plus-excision Cre-Flex and Flp-Flex approaches in HEK293 cells (24). Cre recombinase is the most used and best-understood recombinase (25). Despite this, unexpected Cre-associated behaviours are common in the literature (4, 26–28), and a number of studies have found that Cre can fail to function in genetic circuits implemented in cell-free systems (29), mammalian cells (30) and plant cells (16). In our previous work, a two-input OR gate failed to function with the predicted logic when Cre was expressed (16). Other Cre-related recombinases, such as SCre, VCre, and Vika were also unable to substantially activate output gene expression in a YES gate (16), suggesting that the problem is more widespread than Cre alone, and thereby significantly limiting expansion of the plant recombinase toolkit for circuits. However, a comprehensive understanding of the complex interactions between recombinases, their recognition sites, and target gene elements is still lacking, which confounds predictability in circuit engineering. Here, we sought to address these challenges by characterizing a series of context-dependent transcriptional and recombinase-specific interactions that influence genetic circuit performance in Arabidopsis cells and through this expand the repertoire of genetic tools available for engineering memory circuits in plants.

## Materials and Methods

### Plasmid construction

Plasmid assembly was performed using Golden Gate assembly (31) to create parts (level 0), transcriptional units (level 1), and multigene cassettes (level 2) using plasmid backbones from the Modular Cloning (MoClo) Toolkit (Addgene #1000000044) (32). DNA parts were sourced from the MoClo Plant Parts Kit (Addgene #1000000047) (32) or our past work (16) using the Phytobricks syntax (33) for compatibility and standardization. The nucleotide sequence for synthetic P3-P4 coiled-coil peptides (34), optimized recombinase variant R1-111 and TD1-40 (35) and recombinases KD and B2 (36) were codon-optimized for Arabidopsis expression and synthesized by IDT as gBlock gene fragments. The 5′ UTRs containing recombination sites and *OCS* terminator were synthesized as two separate fragments as IDT gBlocks and assembled into one fragment into a level 0 ligation reaction with *Bpi*I cutting. The truncated SpyCatcher/SpyTag interaction domain (37, 38) was PCR-amplified from the pET28a-SpyCatcher-SnoopCatcher plasmid (Addgene #72324). All plasmids and primer sequences can be found in 10.5281/zenodo.17425901.

### Plant materials and growth conditions

Protoplast experiments were performed using the Arabidopsis Columbia-0 (Col-0) strain and tomato (*Solanum lycopersicum*) cultivar Red Robin. All seeds were stratified at 4 °C for 2-4 days before transfer to the growth chamber and grown at 22 °C under white light and a 16-h day/8-h night diurnal cycle. Arabidopsis and tomato mesophyll protoplasts were generated from plants grown on soil, whereas Arabidopsis root protoplasts were prepared from seedlings grown on half-strength Murashige and Skoog medium agar plates.

### Protoplast transfection

Arabidopsis mesophyll protoplasts were isolated from the leaves of 3-5-week-old plants using the Tape-Arabidopsis Sandwich protocol (39), modified for 96-well plate format (40). Briefly, after enzyme digestion, protoplasts were filtered through a 70 µm strainer, washed with W5 solution and resuspended in MMG solution. Arabidopsis root protoplasts were isolated from seedlings grown on half-strength Murashige and Skoog medium agar plates by slicing the roots and performing enzymatic digestion. Tomato mesophyll protoplasts were isolated by slicing leaves and subjecting them to overnight enzymatic digestion. All post-digestion steps were performed as described above. The cells were counted using a haemocytometer and adjusted to 2 × 10^5^ protoplasts/ml. For transfection, 10,000 protoplasts were mixed with 5 µg of plasmid DNA (in 5 µl) and 55 µl of 40% (w/v) polyethylene glycol. Four, six or eight replicates of each plasmid were assigned to randomized positions on the 96-well plate to minimize positional effects. After 15 minutes, transfection was stopped by adding W5 solution, and protoplasts were incubated for 30 minutes at room temperature before adding WI solution. The transfected protoplasts were incubated under continuous light at 25 °C for 24 hours. Luminescence results from the transfected protoplasts were obtained using the Dual-Luciferase Reporter Assay System (Promega #E1960), and measured on a POLARstar OPTIMA (BMG LABTECH) plate reader.

### Plasmid sequencing

Plasmid assembly was verified by whole-plasmid sequencing on an Illumina MiSeq system. Libraries were prepared through a 5-minute Tn5 tagmentation, where Tn5 transposase, pre-loaded with DNA Illumina adapters, was used to simultaneously linearize and add sequencing adapters to plasmid DNA. The tagmented DNA was then amplified with primers containing Illumina sequencing indexes through ten rounds of PCR with 2x MyTaq (BIO-25041, Bioline). The libraries were pooled together and purified with Serapure beads before sequencing. Reads were assembled *de novo* using Unicycler (version 0.5.0) (41) and aligned to the reference plasmid using Bowtie 2 (version 2.5.4) (42).

### Computational prediction

To investigate potential transcription factors recruited to the recombinase recognition sites, the PlantRegMap (43) Binding Site Prediction tool was used (https://plantregmap.gao-lab.org/binding_site_prediction.php). B3 protein structure prediction was performed by Colabfold (44) via Google Colaboratory to confirm that the proposed truncated protein is of comparable length to Flp recombinase. All structural images used in this study were created in ChimeraX (version 1.10) (45).

### Figure preparation and statistical analysis

The luminescence signals from Rluc and Fluc were generated by the POLARstar OPTIMA plate reader. The output of the circuit was calculated by dividing the output (Rluc) signal by the normalizer (Fluc) signal (Rluc luminescence /Fluc luminescence). Pairwise comparisons were conducted using an F-test followed by Welch’s t-test in R, with p-values adjusted using the Benjamini-Hochberg (BH) procedure. Plots were generated using the ggplot2 package (3.5.1) (46).

## Results

We previously attempted to create an OR gate with Flp and Cre, however this failed to function with the predicted logic in Arabidopsis protoplasts (Fig. 1a) (16). In this circuit, *Renilla luciferase* (*Rluc*) reporter gene expression driven by the constitutive promoter *Act2* was repressed by an *Octopine synthase* (*OCS*) terminator flanked with *FRT* and *loxP* recognition sites for Flp and Cre recombinase binding, respectively, located within the 5’ UTR from the genomic RNA of Tobacco Mosaic Virus (TMV) (Fig. 1a). *Rluc* expression was expected to be restored upon the excision of the *OCS* terminator by either Flp or Cre, however, the *Rluc* reporter failed to be activated whenever the Cre input was present (Fig. 1a) (16). In this study, we showed that a simple one-input switch (YES gate) using Cre only achieved modest activation of the output gene, relative to the Flp recombinase, which led to high output expression (Fig. 1b). Previous work has shown that Cre can bind to the *loxP* site that remains post-recombination and sterically-interfere with transcription (Fig. 1c) (26, 29, 30). To test whether Cre could be inhibiting transcription by binding to the post-recombination target site, we replaced the *loxP* recognition sites with two single mutant target sites, *lox66* and *lox71* (14), that can still efficiently mediate recombination events, but produce a double mutant *lox72* site with low Cre-affinity post-recombination (14) (Fig. 1e). However, we found that the single mutant *lox66/lox71* version of the YES gate only marginally improved circuit performance (Fig. 1d). Therefore, this indicates that the poor performance of these circuits is not only a result of Cre-interference, but potentially also due to additional mechanisms not yet understood.

**Fig. 1:**
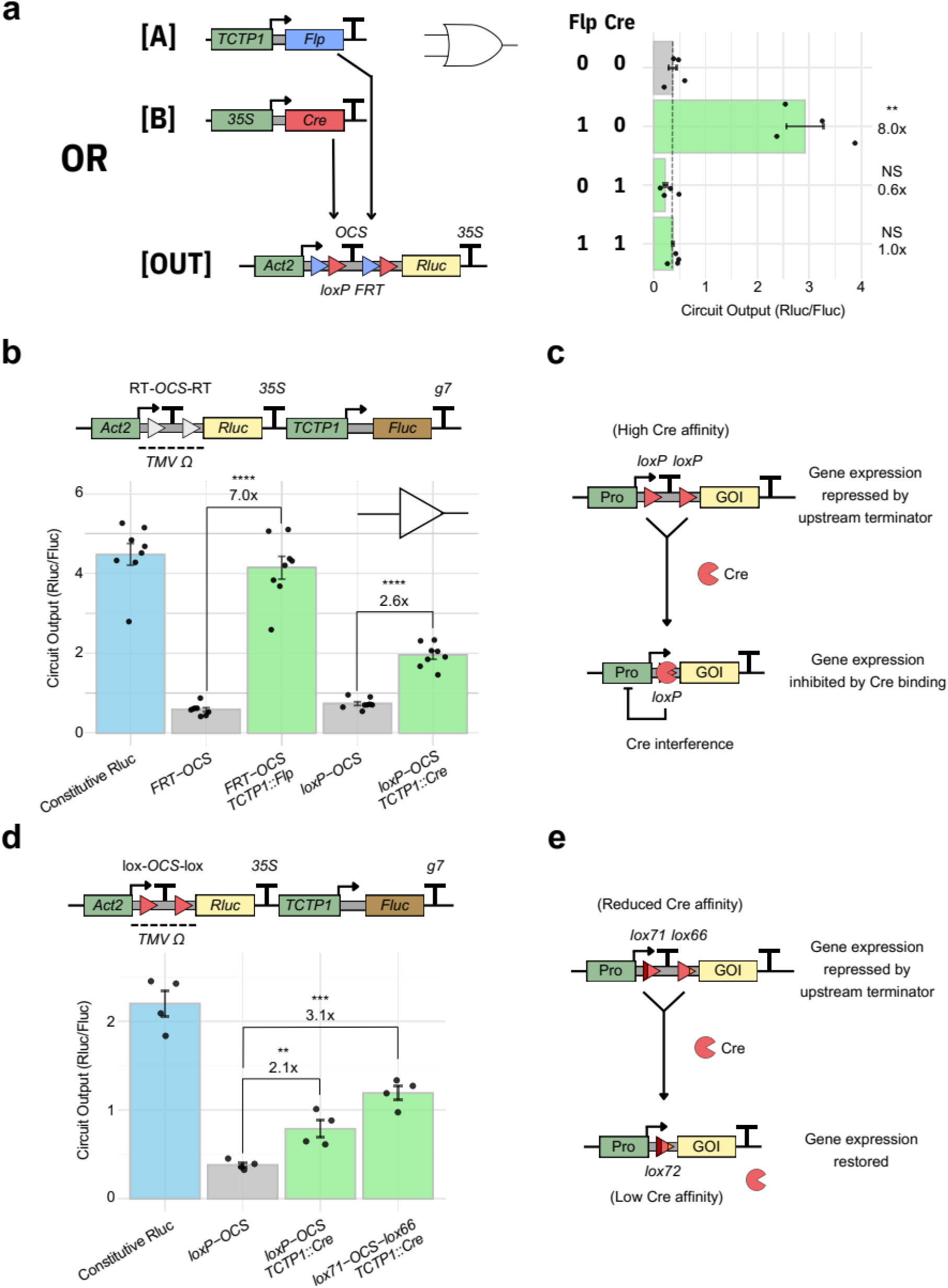
Diagnosing the Cre/*loxP* system in our failed genetic circuit design. **a**. Performance of the original 2-input OR gate with Flp and Cre, measured 24 hours after transfection into Arabidopsis protoplast (*n* = 4). The circuit is activated only in the presence of Flp. Crossbar displays the mean; the blue bar represents the control, the green bar indicates samples expected to be activated, and the gray bar represents samples expected to be repressed. Asterisks indicate a significant difference based on the F-test and Welch’s t-test with Benjamini-Hochberg (BH) adjustment (*P ≤ 0.05, **P ≤ 0.01, ***P ≤ 0.001, ****P ≤ 0.0001; ns, not significant). Adapted from (16). **b**. Comparison of Flp/*FRT* and Cre/*loxP* performance in a one-input YES gate using the *Act2* promoter. Crossbar displays the mean (*n* = 8). Bar colours and asterisks as per panel a. **c**. Schematic of an *Act2* promoter-based 1-input YES gate circuit design with Cre/*loxP*. **d**. Comparison of an *Act2* promoter-based 1-input YES gate using Cre with *loxP* and *lox71*/*66* sites. Crossbar displays the mean (*n* = 4). Bar colours and asterisks as per a. **e**. Schematic of an *Act2* promoter-based 1-input YES gate circuit design with Cre/*lox72*.

To dissect why the Cre-based YES gate did not perform as expected, we examined each component of it in detail. Previous reports of growth inhibition associated with Cre overexpression (47, 48) caused us to consider whether the expression of Cre in the absence of a *loxP* site in the plasmid may alter circuit activity via cellular toxicity. However, we found no such effect (Fig. 2a). We also examined the effect of a single *loxP* site in the 5’ UTR, which replicates the post-recombination state. When Cre was expressed and could bind to the single *loxP* site, a 66% reduction in expression was observed (Fig. 2a), in agreement with the poor activation of the YES gate on-state (Fig. 1d). However, even in the absence of Cre, the output expression was still 33% lower than the control plasmid that had no *loxP* site in the 5’ UTR (Fig. 2a). This indicates that in addition to Cre-interference, the presence of a *loxP* site was also negating the expression of the output reporter driven by the *Act2* promoter (Fig. 2a). This reduction in expression was surprising, given that this is the same position that we had previously inserted other recombinase target sites (*FRT* and *B3RT*) into, with no obvious detriment to circuit performance (16). To determine whether this negation is position-specific, we tested the effect of inserting a single *lox72* site at three different locations within the 5’ UTR. We observed no significant difference in *Rluc* expression between the single *loxP* and *lox72* sites, and none of the alternative placements of *lox72* along the TMV Ω 5’ UTR resulted in higher expression compared to the original (Fig. 2b). This phenomenon of the single *lox* site remaining post-recombination affecting gene expression, which we termed *lox* negation, appears to be a second factor contributing to the failure of our initial circuit design.

**Fig. 2:**
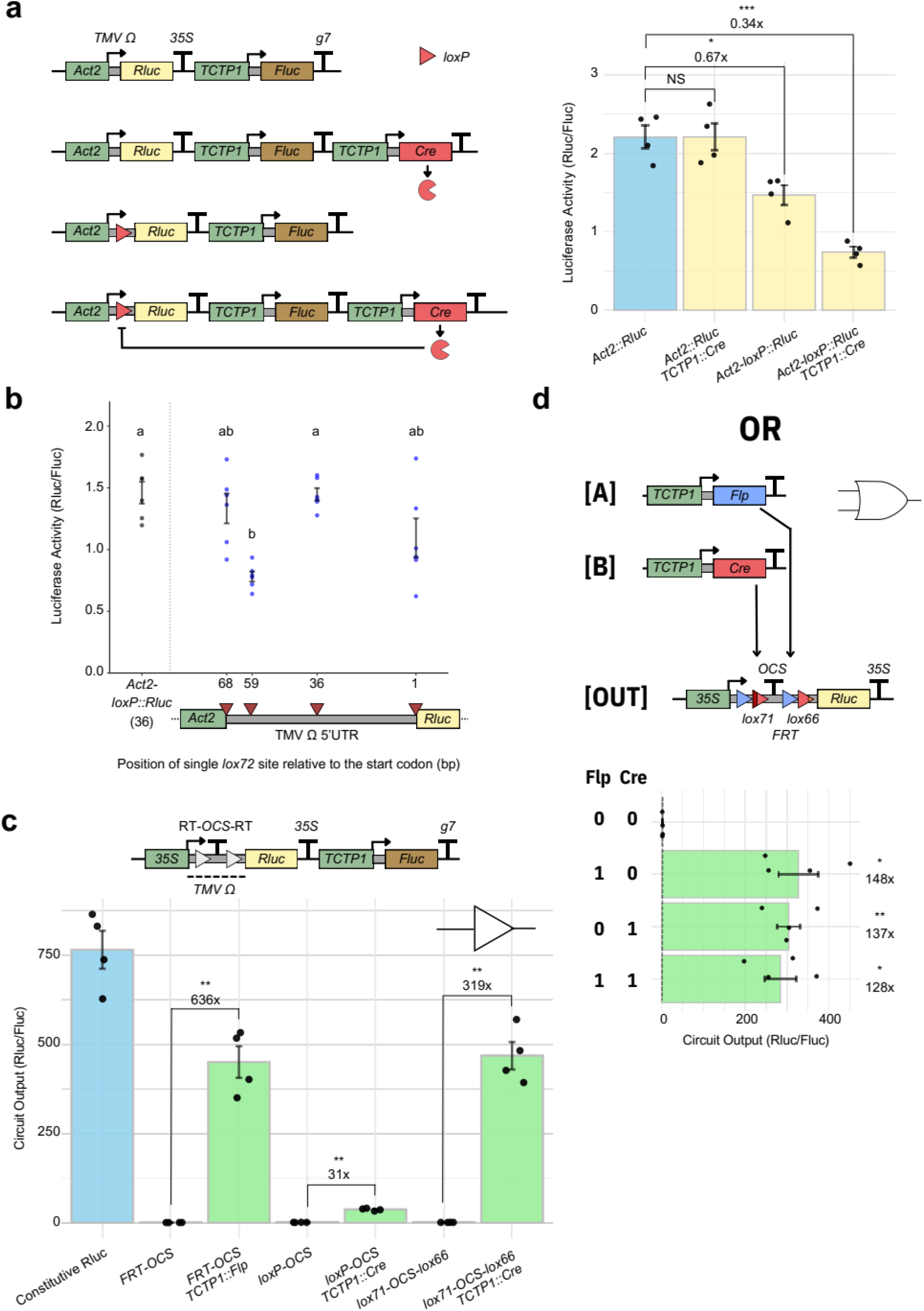
Optimisation of Cre-based logic gates. **a**. Analysis of the *Act2* promoter-based circuit with Cre/*loxP* reveals Cre interference and the intrinsically repressive nature of the *loxP* site. Crossbar displays the mean (*n* = 4); The blue bar represents the control, the yellow bar represents the test sample. Asterisks (****P ≤ 0.0001; *P≤0.01) indicate a significant difference based on the F-test and Welch’s t-test with BH adjustment. **b**. Effect of the *lox72* site in various positions within the 5’UTR driven under the *Act2* promoter. **c**. Comparison of Flp/*FRT*, Cre/*loxP*, and Cre/*lox72* performance in a one-input YES gate using the *35S* promoter. Crossbar displays the mean (n = 4). Colors and asterisks as per Fig. 1. **d**. Optimized two-input OR gate using Cre and Flp, demonstrating high circuit output activity when either input is present. Crossbar displays the mean (*n* = 4). Colors and asterisks as per Fig. 1.

The *Act2* promoter used in our original design is a low/moderate-strength promoter (16, 32). We hypothesized that a more active promoter may be able to counteract the negative effects of the *lox* site and improve reporter gene activation. We therefore selected the Cauliflower Mosaic Virus (CaMV) *35S* promoter, a widely-used promoter in plant biotechnology with a high expression level (49). We replaced the output promoter for the 1-input YES gates with the CaMV *35S* promoter and found that, in the absence of recombinase, *Rluc* expression remains suppressed by the *OCS* terminator located upstream within the TMV Ω 5’ UTR, demonstrating the *OCS* terminator as a robust termination element with low readthrough (Fig. 2c). In the presence of recombinase, we found that all of the YES gates showed significant activation, with a 636-fold increase for Flp and a 319-fold increase for Cre with *lox66/lox71*, respectively (Fig. 2c). This design achieved a greater dynamic range than our previous circuits (16), with >99% repression in the off-state (Fig. 2c). However, when Cre-*loxP* was used, only a 31-fold activation was observed (Fig. 2c), indicating that Cre-interference was still a significant problem when the *35S* promoter was being used. Additionally, we found that the *35S* promoter was resistant to *lox* site negation (Fig. 2c and Supplementary Fig. 1). Given that the issues that were previously observed with Cre-*loxP* (16) could now both be addressed, we decided to re-implement the 2-input OR gate with Cre/Flp as the inputs, taking into account our modifications to circuit design from what we had learned about Cre-*lox* (Fig. 2d). This new version of the OR gate remained repressed in the absence of the recombinase, and demonstrated significant activation in the presence of either or both recombinases (Fig. 2d). Thus, we confirmed that the desired Cre function is restored in the redesigned genetic circuitry, which can be combined with Flp for constructing more complex logic operations, and has a greater dynamic range than our previously published designs.

This curious context-dependent negation of a plant promoter by the presence of a *lox* site led us to investigate if this negation affected other promoters (Fig. 3a). The TMV Ω 5’ UTR with a single *loxP* site was cloned downstream of various constitutive plant promoters: *Act2, TCTP, NOS, AtUbi10*, and *35S*. The *Act2, TCTP*, and *AtUbi10* promoters produced reduced output expression in the presence of a 5’ UTR *loxP* (Fig. 3b-c), whereas *NOS* and CaMV *35S* were unchanged by proximity to *loxP* (Fig. 3c). These data indicate that *loxP* can negate the activity of various plant promoters, but *NOS* and CaMV *35S* are resistant to this effect.

**Fig. 3:**
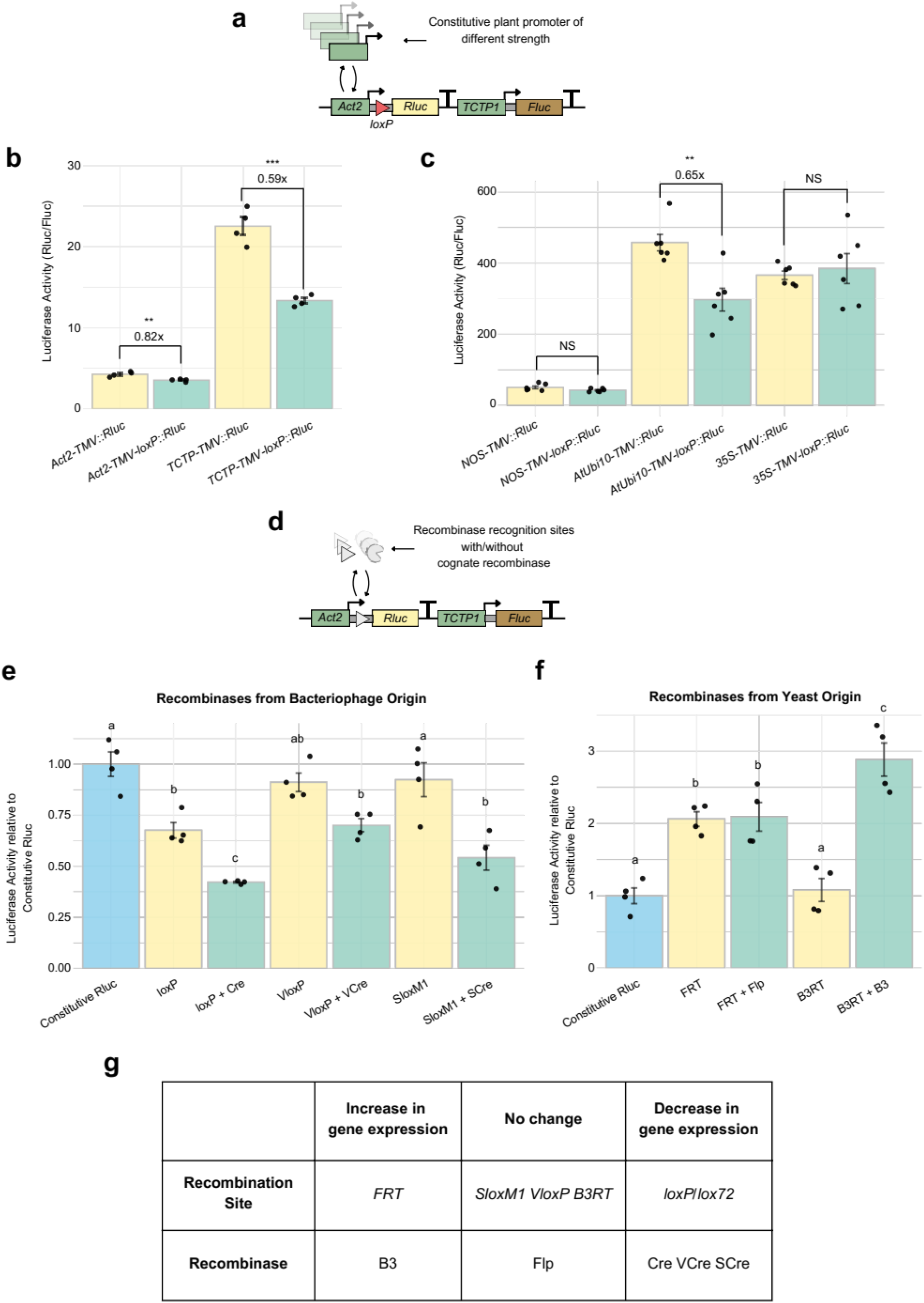
Analysis of the impact of *lox* sites on promoters and the dynamics of different recombinase/recognition site pairs. **a**. Schematics of constructs testing the *loxP* effect on various promoters in plant cells. **b, c**. Impact of *loxP* site on gene expression depending on promoter choice. Crossbar displays the mean (**b**. *n* = 4; **c**. *n* = 6). Yellow bar represents the control sample, while the teal bar represents the sample with recombination site. Asterisks as per Fig. 1. **d**. Schematics of constructs containing different recognition sites and their cognate recombinase. **e, f**. Effect of different recognition sites and recombinase pairs on gene expression. Crossbars display the mean (*n* = 4). Blue bar represents the control sample, yellow bar represents the samples with recombination site, and teal bar represents the samples with recombination site and cognate recombinase. Asterisks as per Fig. 1. **g**. Summary of the effects of different recognition sites and recombinase combinations on gene expression.

Given the impact the *loxP* site can have on expression level, we further examined the single recognition sites for the recombinases Cre, VCre, SCre, Flp, and B3: *loxP, VloxP, SloxM1, FRT*, and *B3RT*, respectively. These sites were inserted into the TMV Ω 5’ UTR in the same position, under the control of the *Act2* promoter (Fig. 3d). This revealed a broad spectrum of interactions between recombinase target sites and their cognate recombinase protein (Fig. 3e-f). Interestingly, while the presence of *loxP* negates gene expression, the *FRT* site caused significant expression enhancement (Fig. 3f). The *VloxP* and *SloxM1* sites had no significant effect on reporter expression, but both VCre and SCre show some degree of recombinase-mediated interference (Fig. 3e), similar to what we had observed for Cre (Fig. 2a). The inclusion of the *B3RT* site did not alter *Rluc* expression, but B3 recombinase augments expression of the output gene, consistent with our previous findings (16) (Fig. 3f). Compared to Flp recombinase, the B3 recombinase has an additional 147 amino acids at the C-terminal (Supplementary Fig. 2a) (35). This is predicted by Alphafold2 (44, 50) to be an intrinsically disordered region (IDR, Supplementary Fig. 2b), which we hypothesize may recruit activator domains to increase expression. Previous work has engineered optimized Flp-like recombinases, R and TD, through the removal of the C-terminal IDR and mutagenesis (35), suggesting that this IDR is unnecessary for recombination. However, removal of this predicted IDR extension from B3 abolished circuit activation, suggesting that the truncated B3 recombinase may no longer catalyze efficient recombination (Supplementary Fig. 2c). Additionally, to determine whether the changes in gene expression from recognition sites alone are due to the presence of motifs for recruiting transcriptional activators and repressors, we scanned each of the recognition sites for transcription factor binding motifs (43). We found that the B3 recognition site may serve as a recognition site for the plant-derived B3 and NAC family transcription factors, although no transcriptional enhancement was observed (Supplementary Table 1). Similarly, *Vlox* recognition sites all contain binding motifs for the AP2 and bZIP family transcriptional factors, while *Slox* recognition sites contain MYB transcription factor binding motifs. No motifs were found in the repressive *loxP* or activating *FRT* sites. Interestingly, we saw no significant change in CaMV *35S* promoter-driven output when these different sites were incorporated into the same position within the TMV Ω 5’ UTR, compared to the native CaMV *35S* promoter (Supplementary Fig. 1). These data show how these seemingly equivalent genetic parts each have their unique characteristics and the importance of fully characterizing each part before complex circuit construction can be undertaken (Fig. 3g).

So far, we have expanded our recombinase toolkit for gene circuits by restoring the function of Cre, but to achieve highly complex logic in plant cells, even more orthogonal components are needed. To achieve this, we tested two homologs of Cre: VCre and SCre, both of which have been shown to function orthogonally to Cre in *E. coli* and mammalian cells (15, 51). Similar to Cre, both VCre and SCre previously demonstrated little or no activation in plant memory circuits (16). We found that they interfere with expression, potentially through tight binding with their native recognition sites (Fig. 3c) (67). We compared the previously characterized *lox66/lox71*-equivalent mutant sites *Vlox43L/R* and *SloxV1L/R* (51), and the mutant site *VloxRSL/R* designed using the previously established DNA specificity profile of VCre (52). Substituting the *VloxP* and *SloxM1* sites with these variants in our *Act2* promoter-based YES gate saw no significant difference between the native/mutant pair *VloxP* and *VloxRS*, as well as *SloxM1* and *SloxV1*, while the *Vlox43* variant only resulted in a slight improvement in activation (Fig. 4a). However, in YES gates based on the CaMV *35S* promoter, all versions of the mutant recognition sites outperformed their native counterparts (Fig. 4b). Furthermore, we tested multiple Flp homologues: KD, B2 and B3, as well as previously optimized variants, R1-111 and TD1-40 (Voziyanova et al. 2016), in *35S* promoter-based YES gates. All designs resulted in significant activation in the presence of their cognate recombinase (Fig. 4c), though the strength of activation varied, likely reflecting the differences in recombinase efficiencies rather than the impact of the recombination site alone. R1-111 and TD1-40 also showed detectable activation when tested in *Act2* promoter-based YES gates (Supplementary Fig. 3). To assess tissue-specific performance, Flp and Cre were tested in Arabidopsis root protoplasts, confirming consistent activation in a distinct cellular context (Supplementary Fig. 4). Additionally, we explored the cross-species functionality of our recombinase circuits by testing the four top-performing recombinases (Flp, Cre, B3 and VCre) in tomato mesophyll protoplasts (Fig. 4d). All four recombinases demonstrated robust activation, indicating that our system retains functionality across species. Together, these findings expand the repertoire of effective recombinases for precise and tunable control of gene expression in plant recombinase-based circuits, thereby advancing the development of more robust and customizable genetic programs.

**Fig. 4:**
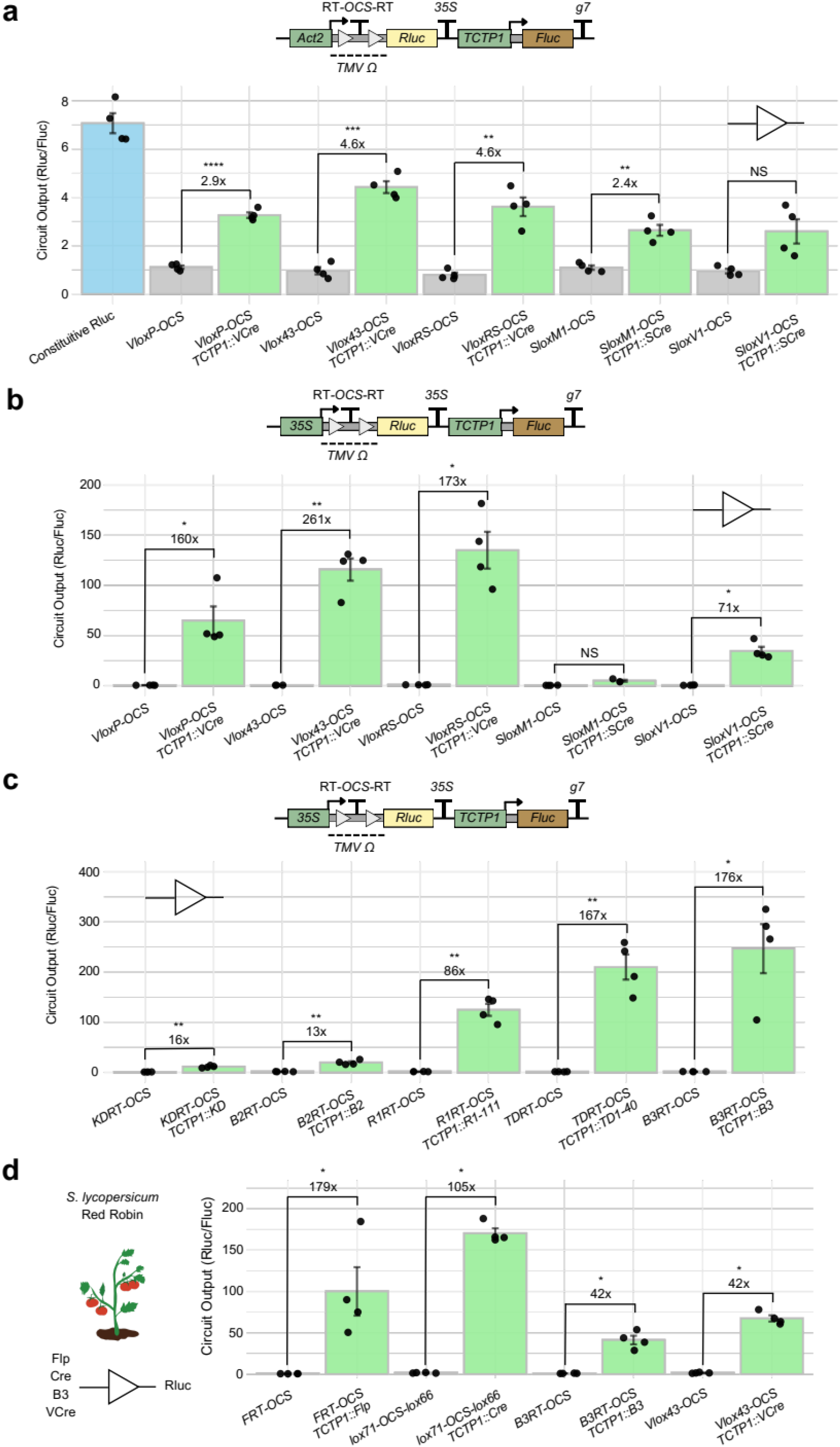
Establishing additional functional recombinases for more complex gene circuits. **a**. Cre homologs VCre and SCre with their native and mutant recognition sites in the context of *Act2* promoter-based 1-input YES gate. Crossbar displays the mean (**a**., **c**., **d**. n = 4, **b**. n = 4 except for the 1-input YES gate with SCre/*SloxM1*, where n = 3), and bar colors and asterisks as per Fig. 1, for all panels. **b**. Cre homologs VCre and SCre with their native and mutant recognition sites in the context of *35S* promoter-based 1-input YES gate. **c**. Flp homologs KD, B2, B3 and engineered variants R1-111, TD1-40 in the context of *35S* promoter-based 1-input YES gate. **d**. Comparison of Flp/*FRT*, Cre/lox72, B3/*B3RT*, and VCre/*Vlox43* performance in a one-input YES gate using the *35S* promoter in tomato mesophyll protoplasts.

Given that we have now expanded the recombinase toolkit for circuits with a range of newly active recombinases, we wanted to demonstrate new complex circuit designs that were not possible to create with our previously limited set of molecular parts for plant memory circuits. We engineered a 3-input AND gate incorporating VCre, Cre, and Flp, achieving 308-fold activation when all three recombinases were present, with no activation in any tested state that lacked any of the three (Fig. 5a). Additionally, the split recombination system involves splitting the recombinase into two inactive halves, which remain inactive unless the split fragments can reconstitute and restore protein function. This system is particularly advantageous in building synchronous logic devices, whereby both inputs must be present at the same time. We have previously shown two-input AND and NAND gates (16) using a functional split-Flp mediated by the C1 homo-dimerisation domain from bacteriophage lambda (53). Here we show similar activation for split-Flp after substituting the C1 domain for a truncated version of SpyCatcher/SpyTag from *Streptococcus pyogenes* (37, 38), and a synthetic pair of coiled-coil dimers (P3-P4) (34) (Supplementary Fig. 5). Thus, we took advantage of the known split-Cre and split-VCre sites (54) and fused them with the P3/P4 coiled-coil dimers and the SpyCatcher/SpyTag, respectively. For both split-recombinases, we observed no activation of the repressed *Rluc* gene in the absence of the recombinase or when only one fragment was present (Fig. 5b-c). Only when both fragments were present was a high level of *Rluc* activation detected in Arabidopsis (Fig. 5b-c) and tomato (Supplementary Fig. 6). Expanding on the split recombinase system and our functional 3-input AND gate, we split each of the VCre, Cre, and Flp recombinases into two fragments, fusing them with SpyCatcher/SpyTag, P3-P4 coiled-coil, and C1 domains, respectively. Despite this added complexity, the system remained functional, exhibiting strong (159-fold) activation when all six components were present (Fig. 5d). By expanding effective split recombination systems in plants and demonstrating their capacity for multi-input logic operations, these results open new possibilities for constructing increasingly sophisticated genetic circuits.

**Fig. 5:**
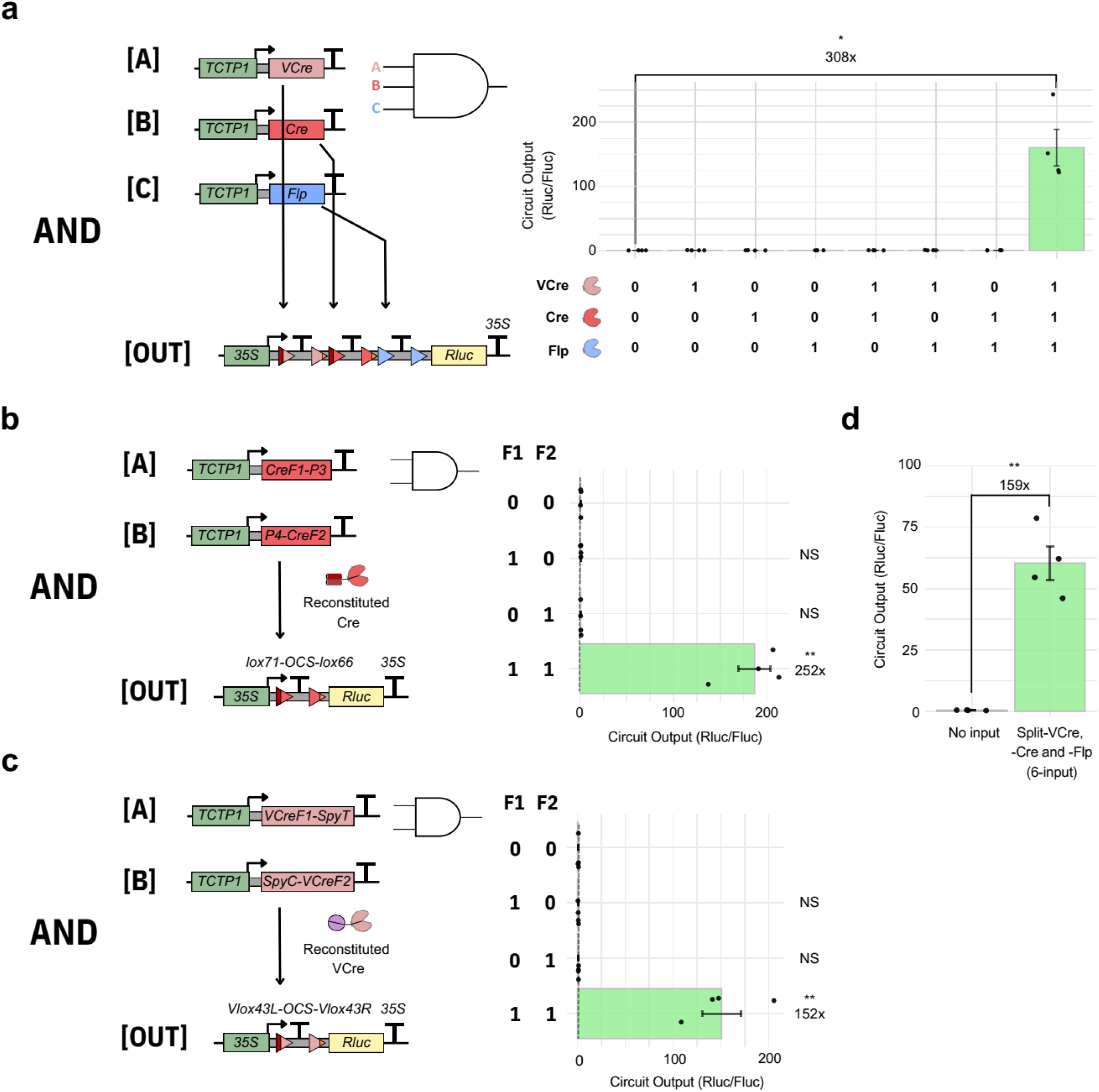
Complex genetic circuit design with expanded recombinases. **a**. Optimized three-input AND gate using VCre, Cre, and Flp, demonstrating high circuit output activity when all inputs are present. Crossbar displays the mean (n = 4) and bar colors and asterisks as per Fig. 1, for all panels. **b**. Two-input split-AND gate with split-Cre, fused to P3-P4 coiled coil domains. **c**. Two-input split-AND gate with split-VCre, fused to the SpyTag/SpyCatcher domains. **d**. Performance of the AND gate design with split-VCre, -Cre and -Flp.

## Discussion

Our ability to generate user-defined changes in plant traits with synthetic gene circuits is limited by the molecular toolkit that is available to us to reprogram plant gene expression. In this work, we expand the tools available for memory circuits by investigating the underlying cause behind the lack of predicted activity of circuits using Cre, VCre and SCre (16). It has been established that post-recombination Cre can recognize and bind to this residual *loxP* site and interfere with gene expression in a manner akin to CRISPR-interference, in which dCas9 represses targeted genes by obstructing RNA polymerase transcribing through the region (55, 56). By modifying the affinity of Cre for the post-recombination target site with mutations to generate a *lox72* site, some activity could be restored (29, 30). However, we found that circuit activation was still much less than predicted, leading us to discover that *loxP* had a repressive effect when placed in a plant 5’ UTR, downstream of a range of different promoters. Among the constitutive promoters tested, only the strong viral *35S* promoter and the moderate *NOS* promoter were resistant to this repressive property of *loxP*. Furthermore, we were able to identify changes to the target sites of SCre and VCre that allow for improved circuit activation with these recombinases. These optimized recombinases were further engineered in their split forms to enable the construction of multi-input logic gates. Taken together, we have been able to significantly increase the toolkit available for building effective plant memory circuits, and outline pitfalls and accompanying solutions to issues that can affect any expression platform that utilises recombinases, regardless of the species.

The expanded repertoire of recombinases for plant memory circuits established in this study not only facilitates the construction of more complex logic operations but also allows for the precise control of multiple orthogonal pathways within the same cell. These new tools can be integrated with the existing recombinase toolbox to engineer more sophisticated recombinase-based circuit architecture for complex cellular logic. This approach will enable improved spatiotemporal regulation of gene expression in plants, mediated by the simultaneous activity of multiple promoters, which can be especially valuable for engineering crops with advantageous traits. For instance, a two-input AND gate could combine a development stage-specific promoter with a chemical-responsive promoter to activate a gene encoding a transcriptional factor that triggers the accumulation of health-promoting metabolites only when the plant reaches a specific development stage and the chemical signal is present. This strategy allows for precise gene expression control, reducing the risk of abnormal growth phenotypes due to broad expression domains associated with constitutive expression (57). Thus, expanding the recombinase toolkit advances synthetic biology in plant engineering and offers new opportunities for fine-tuning trait development.

Beyond the expansion of recombinase-based genetic circuits in plants, our study highlights the unique set of challenges when attempting to build synthetic gene circuits in any biological system. Unlike its electrical counterpart, incomplete knowledge of genetic interactions, variation between species, and unknown cellular characteristics mean that even a simple genetic circuit can exhibit unanticipated behaviors. Prior works on recombinases have looked in-depth at altering the recombination efficiency (58, 59), enhancing site-specific accuracy (60), engineering recombinases for novel site specificity (61), and the expansion of orthogonal recognition sites (62). Yet, the effect of the post-recombination target site scar has not been well analysed. While the impact of this site’s interaction likely varies across organisms, our findings confirm that these short residual sequences have the potential to influence gene expression. Our initial *loxP* placement is located within the central region of the TMV Ω 5’ UTR, which contains a poly(CAA) region responsible for translation enhancement through the recruitment of heat shock protein HSP101 (63). While we hypothesized that the presence of the recombination site may disrupt the region’s ability to serve as the binding site, the alternative sites selected specifically to avoid the poly(CAA) region did not improve gene expression (Fig. 2b). Furthermore, the introduced recognition site sequences may serve as sites for transcriptional activators or repressors. We also discovered that this recombination site-dependent effect persists across other studies and can reduce the expression of proximal genes (64), or completely abolish downstream gene expression (65). Given that recombinase recognition sites are often palindromic, except for the core sequences that determine directionality, it is plausible that the resulting mRNA could form hairpin structures, potentially destabilizing the transcript and affecting the output luciferase level. Thus, the presence of the residual recombination site post-recombination is capable of modulating output activity, a factor that needs to be taken into consideration when fine-tuning genetic circuits with increasing complexity.

We have demonstrated that the presence of the cognate recombinase on the recombination site in the aftermath of recombination needs to be taken into account. The ability for recombinase to affect gene expression post-recombination is evident in that Cre-interference can drastically diminish the output (Fig. 1b & 2c) (29, 30). Similar effects have been observed with serine integrases, where high recombinase-attachment site affinity was shown to suppress downstream gene expression (66). Fortunately, because the overexpression of Cre causes undesired effects (47, 67), most approaches prefer to express Cre through an inducible system (54, 68), which minimizes the risk of Cre-interference, but this can be a problem if Cre is maintained after recombination has taken place (15, 16). Additionally, as the promoter of choice for driving circuit output in literature implementing recombinase is typically strong, such as human *elongation factor-1 alpha* (*EF1α*) (15) in mammalian cells and CaMV *35S* promoter in plants (69), the activation generated, albeit at a magnitude lower than what could be accomplished, may be enough for most applications. The interference caused by Cre may also have a drastic effect on weaker promoters, as was the case for the *Act2* promoter (16). We saw no such interference in the yeast-derived Flp recombinase, likely due to the lower affinity Flp has to the *FRT* site in comparison to Cre/*loxP* (70). Intriguingly, we saw an enhancement in transcriptional activity in the case of B3 binding, suggesting that B3 may act as a transcriptional activator, in addition to a recombinase, at least in plant cells. The IDR in the C-terminus of B3 may play a role in recruiting activator domains to the region, which we attempted to diagnose by removing the extension compared to Flp (Supplementary Fig. 2a). However, the removal of the region abolished recombination activity from B3 (Supplementary Fig. 2c), although subsequent mutagenesis may be possible in yielding a truncated B3 recombinase.

Despite the apparent similarity in the mechanism of action of tyrosine recombinases (71) and length of their target site (33-34 bp), most recombinase-target site pairs examined in this study exhibited unique behaviors, even when placed in the same genetic context (Fig. 3e-f). Cre, VCre and SCre proteins all interfere with expression when able to bind to their target site (Fig. 3e), but only *loxP* target site negates promoter activity (Fig. 2a, 3b, 3e), whereas *VloxP* and *SloxM1* have no effect on expression (Fig. 3e). Interestingly, the binding of the B3 recombinase to its target site augments output gene expression (Fig. 3f), which could be a useful tool for further boosting target expression. However, the *B3RT* site appears inert and does not alter expression without the presence of the B3 recombinase. Conversely, Flp recombinase binding does not affect transcription, yet the *FRT* site independently enhances expression of the output gene (Fig. 3e). Collectively, these findings highlight that recombinases and their target sites cannot be used as interchangeable genetic parts. Instead, a detailed analysis of each component within a gene circuit is essential for accurately predicting circuit activity, and we believe this is a valuable lesson for any project within synthetic biology.

Overall, our findings demonstrate that while the site-specific recombinases are powerful tools for constructing memory-based logic gates with decision-making capabilities, the interactions between genetic components must be carefully considered to ensure reliable circuit performance. This study not only expands the toolkit for designing memory circuits in plants, but also highlights the need for a deeper understanding of how recombinase activity, recognition sites, and surrounding genetic elements interact. This work offers lessons for how to best design applications of Cre and related recombinases that we believe will be valuable in diverse organisms beyond plants. By addressing these dynamic interactions, our work serves as both an expansion of plant-based tools for recombinase-based gene circuits and a cautionary tale highlighting the importance of understanding context-specific behavior when designing synthetic gene circuits.

## Acknowledgements

The Plasmid-seq data were generated on instrumentation supported by the Australian Cancer Research Foundation Centre for Advanced Cancer Genomics and Genomics WA.

## Author Contribution

P.G., J.P.B.L. and R.L. conceived of the project, designed the experiments and analyses, and wrote the manuscript. P.G. conducted the experiments with assistance from J.P.B.L., A.K., F.L., J.Z., G.H., H.J., and C.P. All authors approved of and contributed to the final version of the manuscript.

