## Supplementary figures for "CREation of an expanded plant memory gene circuit toolkit"

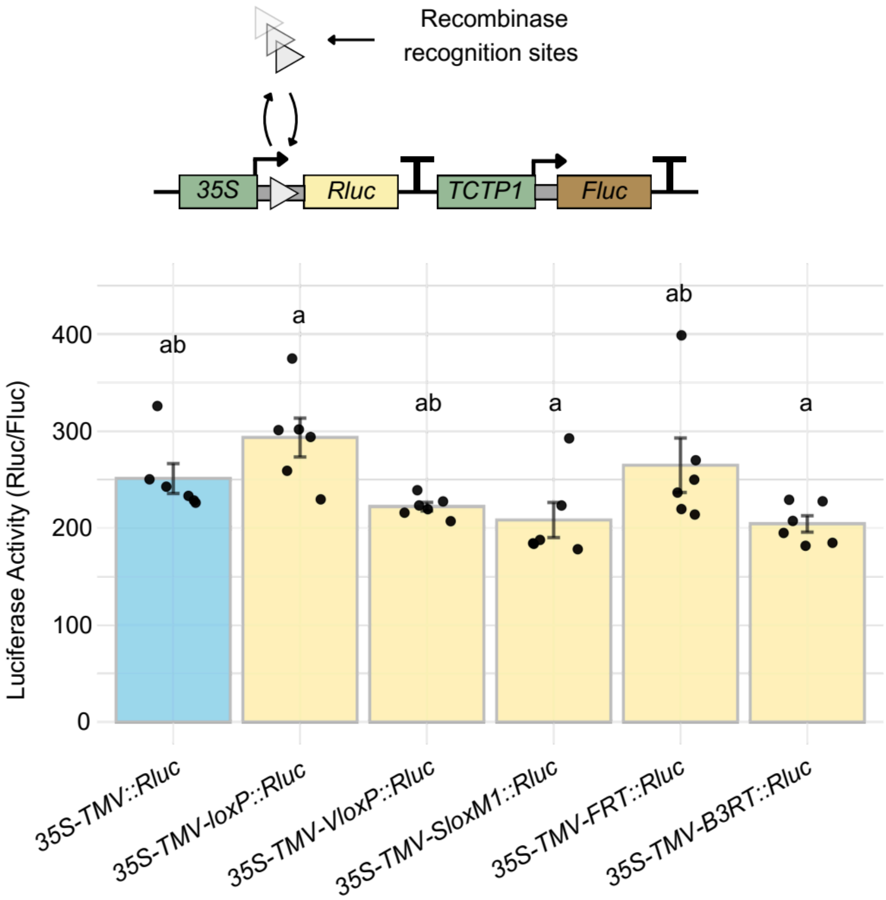


**Supplementary Fig. 1: Analysis of the dynamics of different recognition sites under the *35S* promoter**.

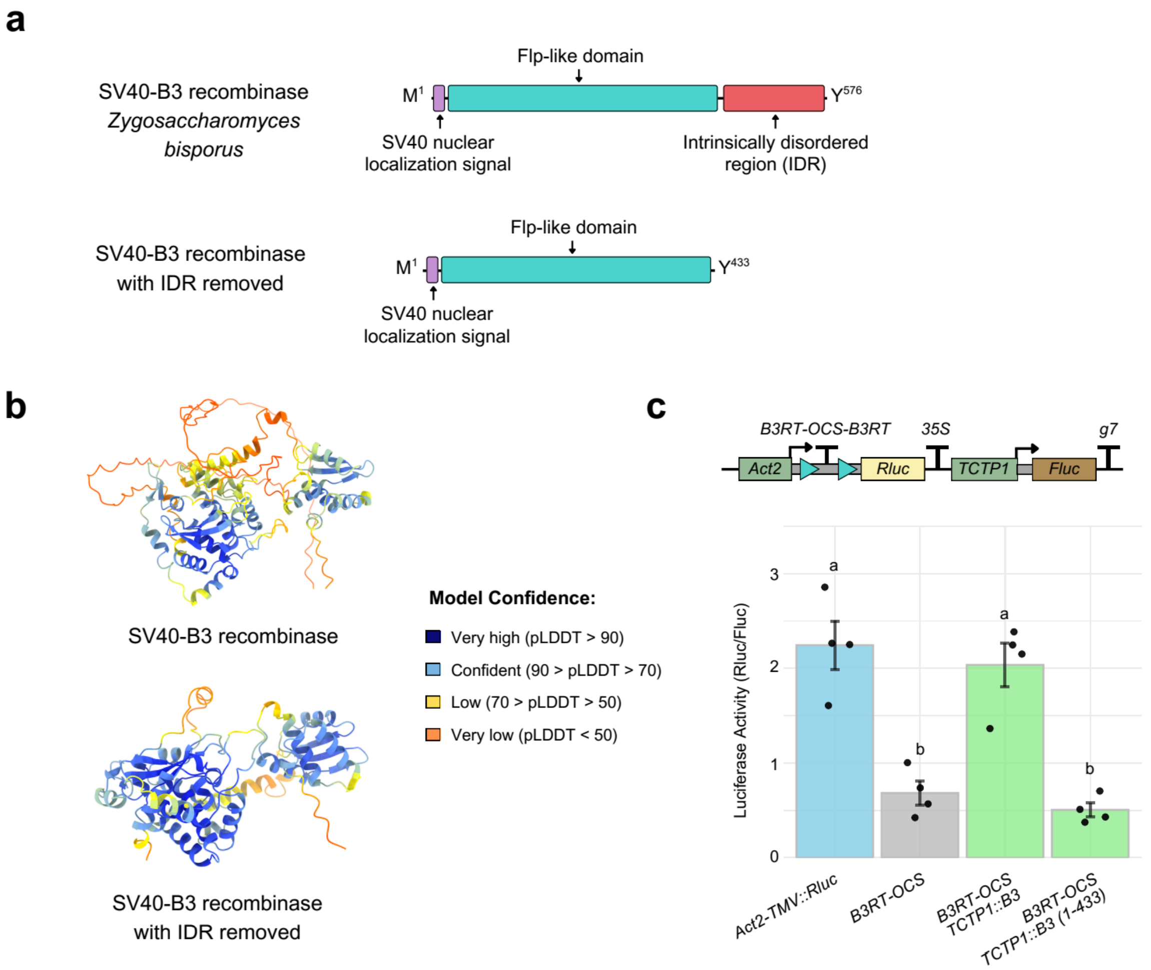


**Supplementary Fig. 2: Truncated B3 recombinase with the removal of the intrinsically disordered region (IDR)**. **a.** Line diagram of the B3 recombinase, where different colors indicate the different domains. **b.** Alphafold prediction of B3 recombinase and its truncated variant with IDR removed. Structure was generated with ChimeraX (Meng et al. 2023). **c.** Comparison of an *Act2* promoter-based 1-input YES gate using full-length and truncated [B3 (1-433)] B3 recombinases.

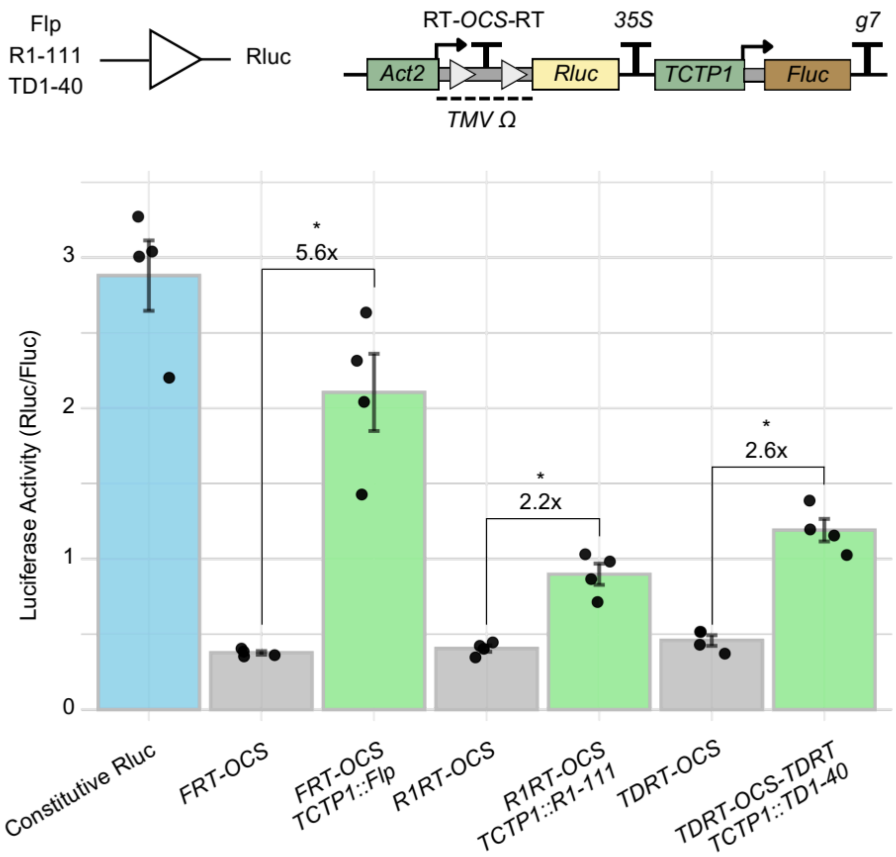


**Supplementary Fig. 3: Comparison of Flp/FRT, R1RT/R1-111, and TDRT/TD1-40 performance in an *Act2* promoter-based 1-input YES gate.**

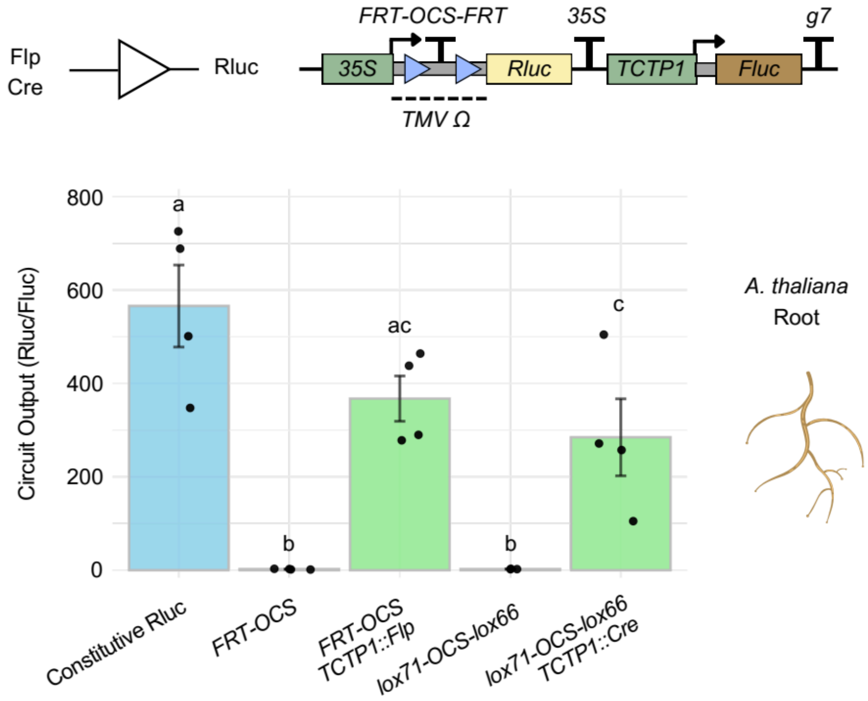


**Supplementary Fig. 4: Comparison of Flp/FRT and Cre/lox72 performance in a one-input YES gate using the *35S* promoter in Arabidopsis root protoplast.**

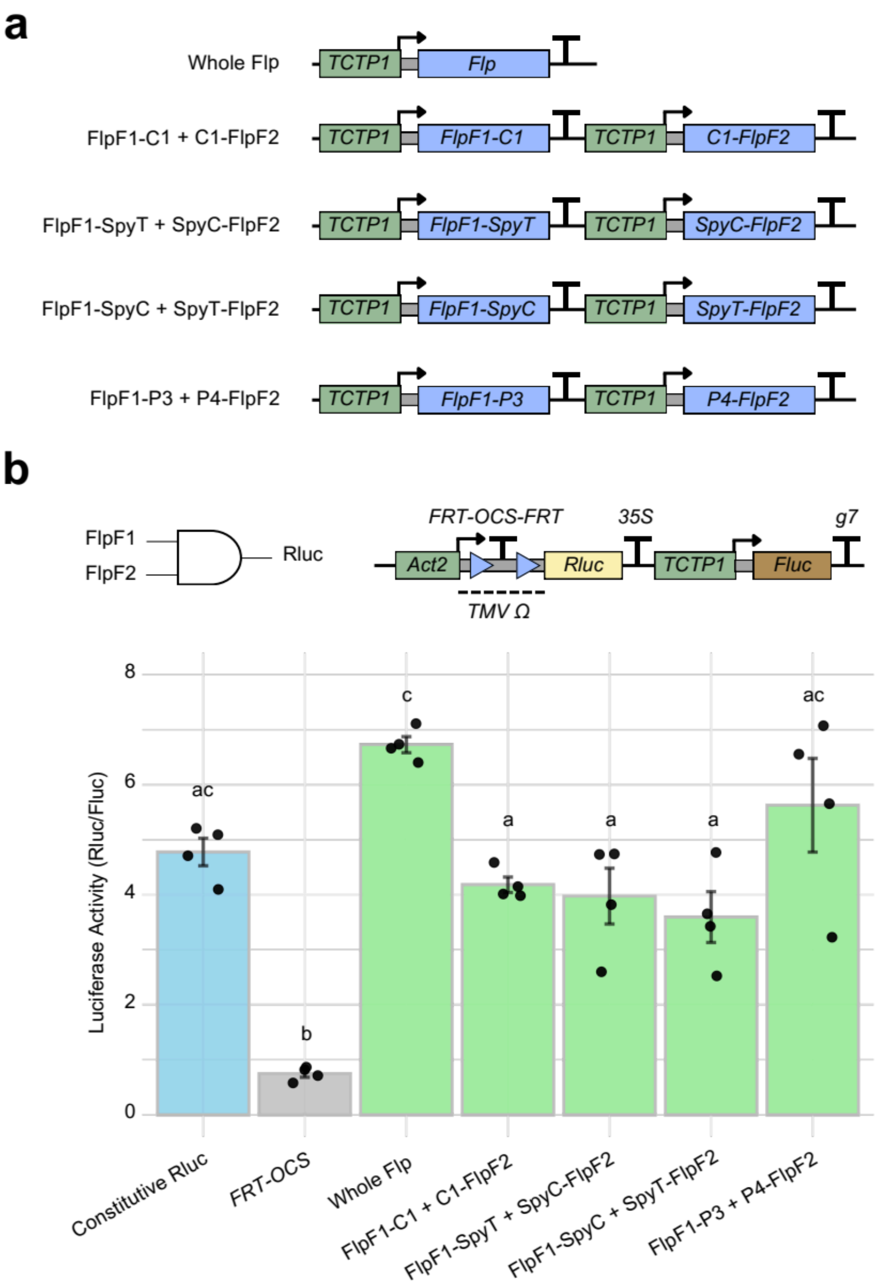


**Supplementary Fig. 5: Split-Flp with different dimerization domains**. **a.** Schematics of whole Flp and split-Flp fused with different dimerization domains. **b.** Comparison of an *Act2* promoter-based 2-input AND gate using split-Flp fused with different bioconjugation domains.

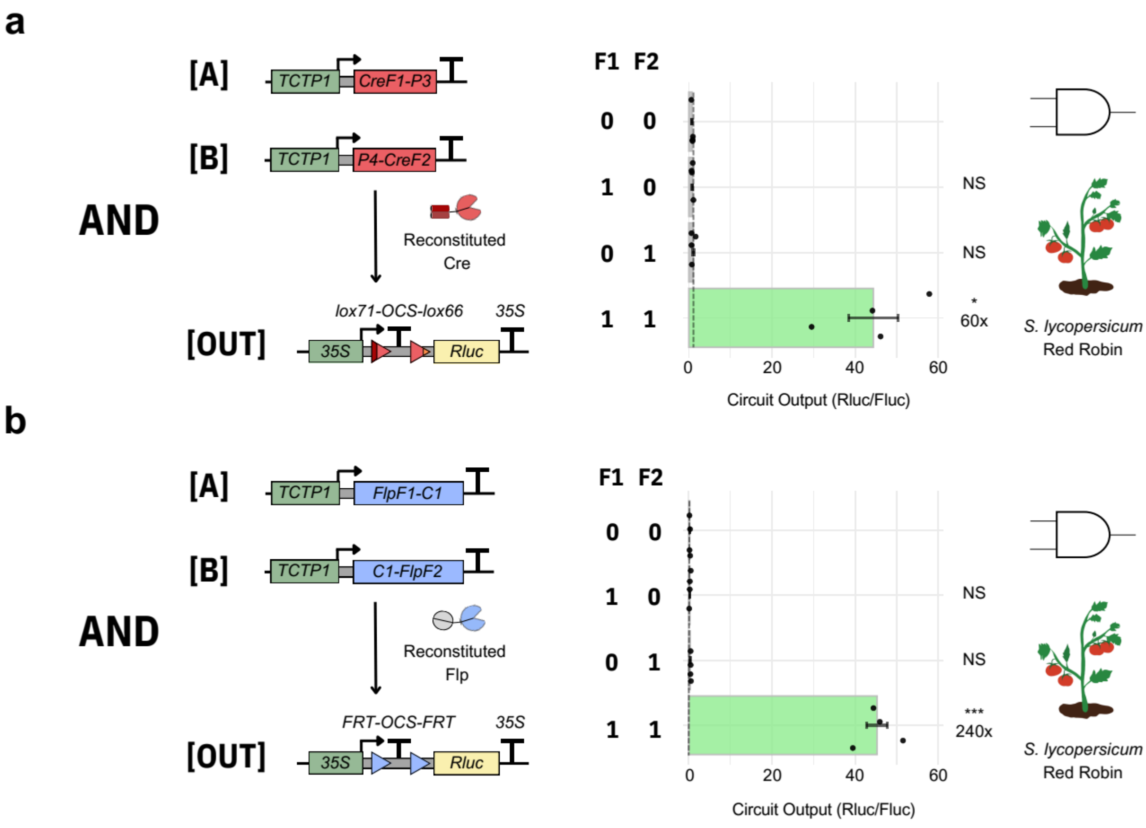


**Supplementary Fig. 6: Complex genetic circuit designs in tomato mesophyll protoplast. a.** Two-input split-AND gate with split-Cre, fused to P3-P4 coiled coil domains. **b.** Two-input split-AND gate with split-Flp, fused to C1-C1 homodimerization domains.
